# Systems-level reconstruction of kinase phosphosignaling networks regulating endothelial barrier integrity using temporal data

**DOI:** 10.1101/2024.08.01.606198

**Authors:** Ling Wei, John D. Aitchison, Alexis Kaushansky, Fred D. Mast

## Abstract

Phosphosignaling networks control cellular processes. We built kinase-mediated regulatory networks elicited by thrombin stimulation of brain endothelial cells using two computational strategies: Temporal Pathway Synthesizer (TPS), which uses phosphoproteomics data as input, and Temporally REsolved KInase Network Generation (TREKING), which uses kinase inhibitor screens. TPS and TREKING predicted overlapping barrier-regulatory kinases connected with unique network topology. Each strategy effectively describes regulatory signaling networks and is broadly applicable across biological systems.

---

Protein kinases regulate numerous cellular processes through phosphorylation of target proteins. These series of phosphosignaling events are triggered in response to environmental changes and propagate within a network structure. The function of kinases in cell signaling and regulation of cellular phenotypes has been intensively explored in different contexts, including in mediating brain endothelial barrier integrity ^1^. The integrity of the blood-brain barrier is essential for maintaining the homeostasis of the central nervous system ^2^. The permeability of brain endothelium is tightly controlled by cell-cell junctions and focal adhesions, whose functions are heavily regulated by kinases ^1–3^. Therefore, understanding the network-level connectivity of these barrier-regulatory kinases and their temporal dynamics is critical for developing targeted therapeutic strategies for treating vascular leakage.

In recent years, computational tools and modeling approaches have been advanced to facilitate reconstructing high-confident, systems-level protein signaling networks based on proteomics data and prior knowledgebases (Table 1), but they often fall short of providing detailed mechanistic understanding and/or fine temporal information on cell signaling and their function in mediating cellular phenotypes. For example, Babur *et al.* developed a pathway extraction method that maps high-throughput proteomics data to the Pathway Commons database. They used a graphical pattern search framework to identify the protein-protein interactions (PPIs) that explain correlated changes in proteomics profiles ^4^. Network inference methods like dynamic Bayesian networks have been implemented on time course phosphorylation data to infer protein signaling network structure specific to biological context without relying on user-defined parameters ^5–7^. Ordinary differential equation (ODE) models are kinetic models that have been conventionally applied to analyze biochemical reaction networks, which can capture the mechanistic details of biochemical processes and predict the temporal evolution of variables that cannot be directly measured ^8^. Unfortunately, building robust medium- or large-scale ODE models is challenging due to a lack of detailed mechanistic description of the biological system and insufficient experimental data to parameterize the models ^8,9^. Unbiased network reconstruction has benefitted from the development of logic modeling, which employs either discrete logic models or continuous ODEs transformed from discrete logic models, followed by optimization to fit experimental data, to generate predictive models that describe context-specific signaling within the generic protein signaling networks ^10,11^.

**Table 1.**
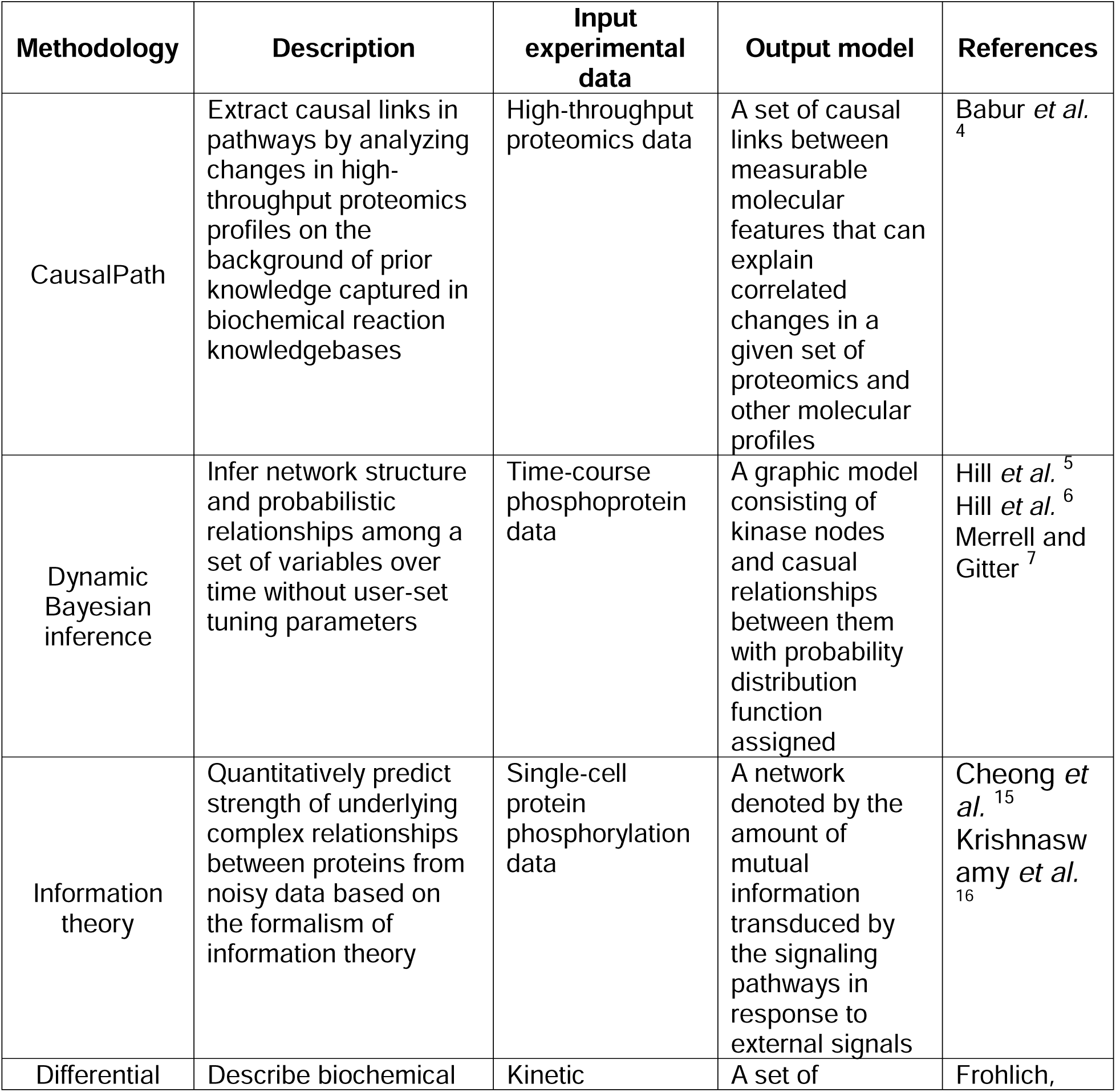
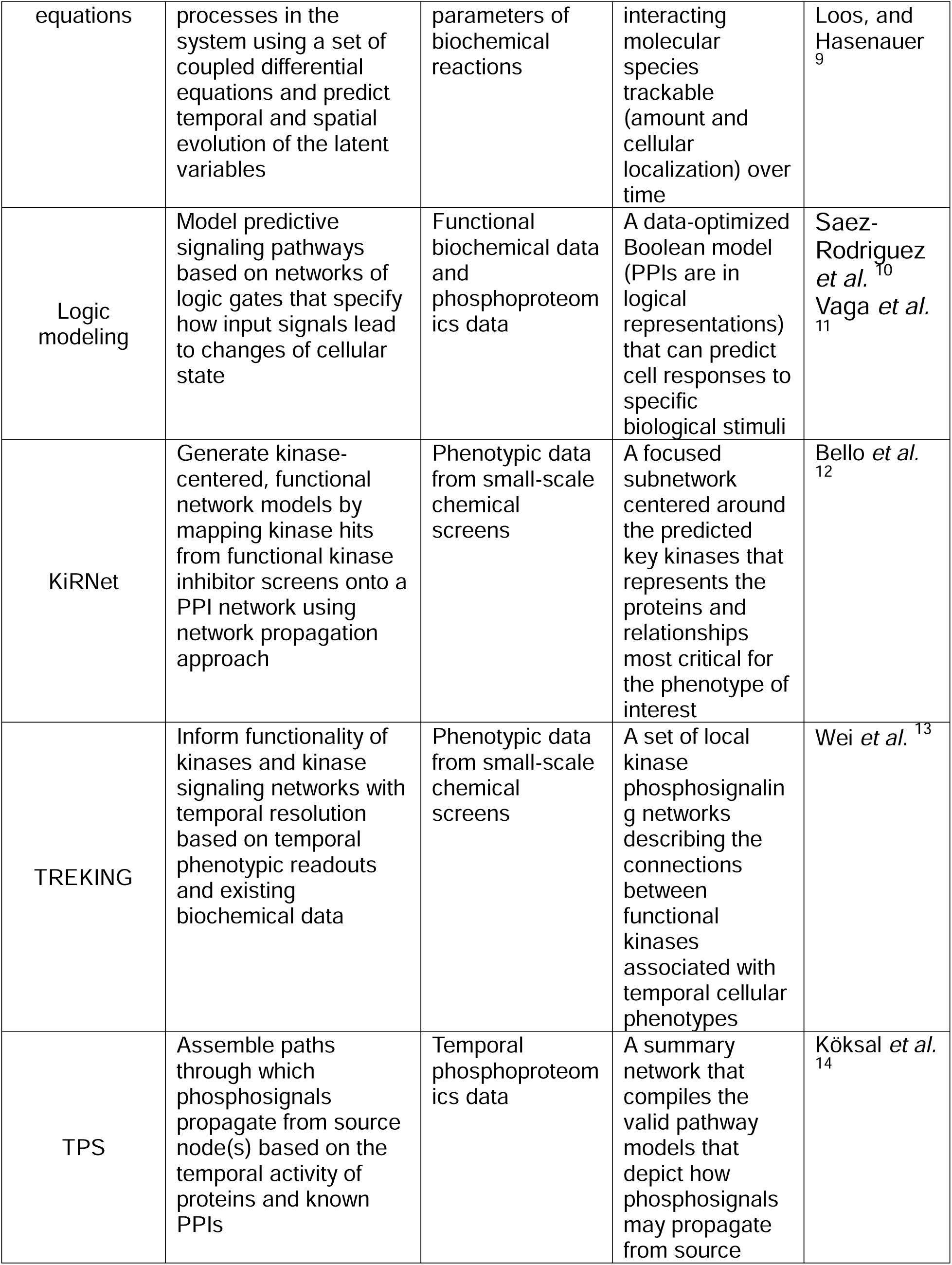
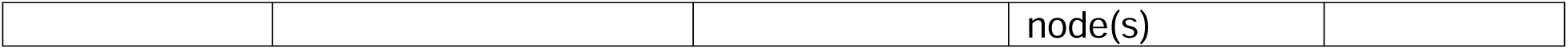
List of computational approaches for building and analyzing protein phosphosignaling networks.

Besides building signaling networks from phosphoproteomics data, which are typically collected at relatively sparse time points, network generation pipelines can use functional screens with fine temporal resolution as input along with known PPIs ^12,13^ (Table 1). For example, Bello *et al.* extended the kinase regression (KiR) approach that was initially designed to identify key kinase regulators of a particular cellular phenotype from quantitative drug screens. Their KiRNet method maps KiR hits onto existing PPI networks using network propagation to create kinase-centered, functional network models associated with the phenotype of interest ^12^. We recently developed a novel approach called TREKING that integrates temporal drug screen data with a known kinase-substrate phosphorylation database to infer functional kinase signaling networks with temporal resolution ^13^.

Köksal *et al.* have also developed a network inference algorithm specifically to model time course data ^14^. The Temporal Pathway Synthesizer (TPS) uses time-resolved phosphoproteomics data as input ^14^. TPS assembles signaling pathways by first extracting a subnetwork that connects the phosphorylated proteins to the source node(s) from a background PPI network. It then uses the temporal phosphorylation data to systematically examine all directed, signed edges in the subnetwork to determine how signals propagate from source node(s). Finally, TPS consolidates all valid paths into a unified network that represents the pathway structures facilitating phosphosignal propagation.

Phosphoproteomics-based kinase signaling networks offer a distinct perspective from those derived from functional assays. While phosphoproteomics help elucidate how phosphosignals propagate within phosphosignaling networks, functional assays can link kinase activity directly to cellular phenotypes. Comparing these two network reconstruction approaches enhances our understanding and interpretation of phosphosignaling events crucial to the phenotype of interest.

Here, we reconstructed a phosphosignaling network regulating thrombin-induced endothelial barrier disruption and recovery using the TPS algorithm (Fig. 1). We then compared this network with a TREKING network we previously assembled using kinase inhibitor screens in the same cellular background. The input time series data for TPS were collected from western blot experiments as described in ^13^. Briefly, human brain microvascular endothelial cells (HBMECs) were treated with thrombin and the phosphorylation status on the activation sites of 25 kinases, and 3 non-kinase proteins known to regulate barrier function, were collected at nine time points across a 6-hour time window after thrombin treatment. Data on 17 kinases have been published previously ^13^, and data generated in this study on the remaining 8 kinases and 3 non-kinase proteins are available in Supplementary Data S1. The input network that connects phosphorated proteins to the source node, proteinase-activated receptor 1 (F2R/PAR1), was generated using Omics Integrator ^17,18^ (Supplementary Data S2). The input files for TPS were prepared according to ^14^ and are described briefly in Methods.

**Fig. 1.**
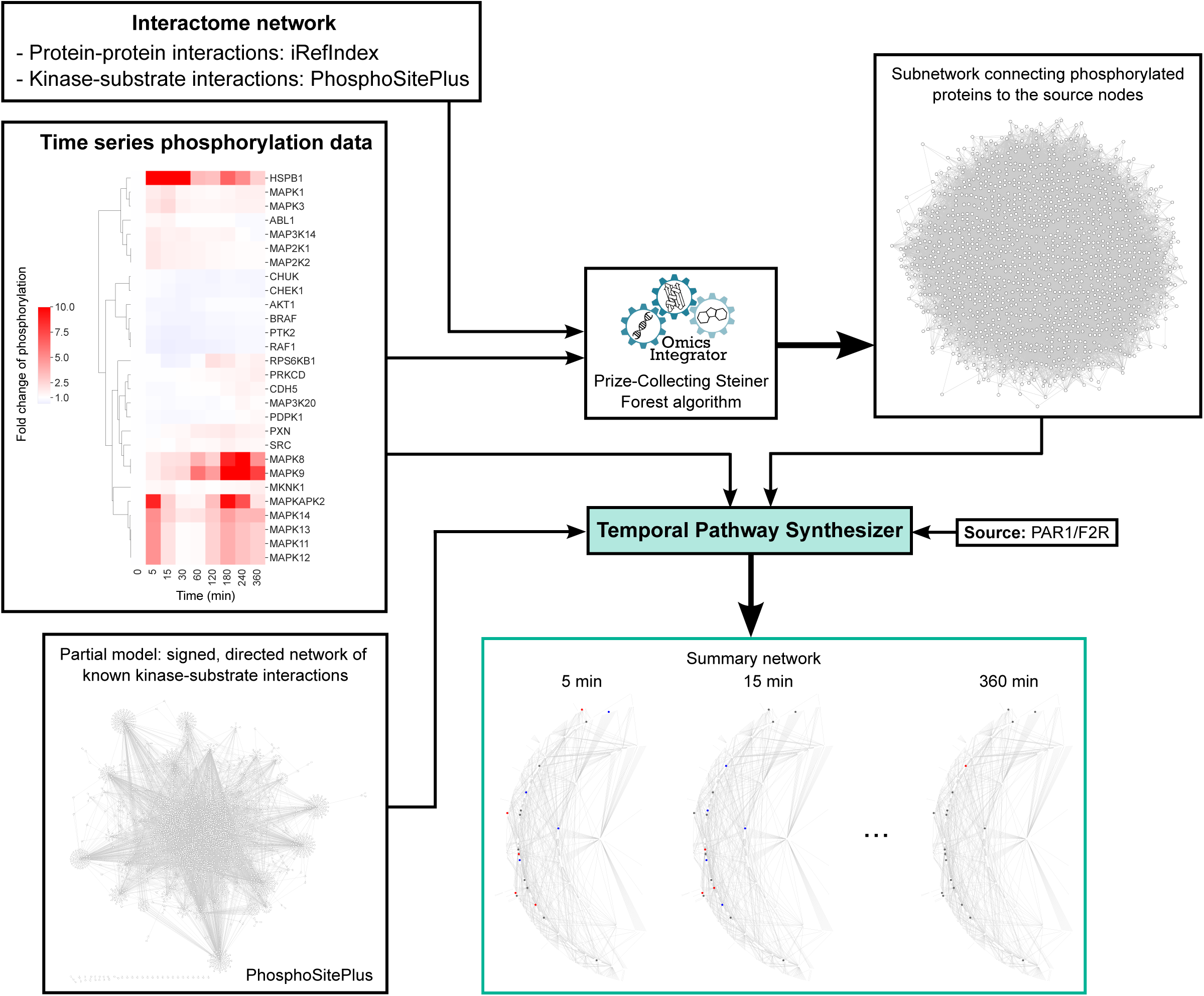
Workflow for reconstructing a kinase-mediated phosphosignaling network controlling brain endothelial barrier integrity after thrombin stimulation using TPS. HBMECs were treated with thrombin for 0, 5, 15, 30, 60, 120, 180, 240 or 360 minutes. Phosphorylation levels (at activation sites) of 28 proteins (25 kinases and 3 non-kinase proteins) were evaluated by western blot and quantified by fold change from the basal level (media only) after normalization to GAPDH. The interactome network was compiled by combining human PPIs from iRefIndex and kinase-substrate phosphorylation interactions from PhosphoSitePlus. An undirected subnetwork was generated using the Omics Integrator implementation of the Prize-Collecting Steiner Forest algorithm, incorporating the time series phosphorylation data and the interactome network. The subnetwork, kinase-substrate phosphorylation interactions, and time series phosphorylation data were input into the TPS algorithm, with PAR1/F2R as the source node. TPS outputs included 1) the kinase-mediated signaling network (directed and signed) that controls protein phosphorylation in HBMECs after thrombin treatment, and 2) activity of each protein in the subnetwork at the corresponding time points.

The TPS-generated network resulted in 966 directed edges, with only five signed, directed edges due to the sparsity of the input time series data (Fig. 2a, Supplementary Data S3). The network recapitulates interactions between extracellular signal-regulated kinases (ERKs; MAPK1/ERK2 and MAPK3/ERK1) and β-arrestins (ARRB1 and ARRB2), key components of canonical thrombin signaling through PARs (Reactome pathway R-HSA-456926: thrombin signaling through proteinase activated receptors (PARs), Supplementary Data S5). The model predicts that both ABL proto-oncogene 1 non-receptor tyrosine kinase (ABL1/c-Abl) and mitogen-activated protein kinase 8 (MAPK8/JNK1) activate paxillin (PXN), consistent with previous findings ^19,20^. However, the model’s prediction that protein tyrosine kinase 2 (PTK2/FAK) inhibits PXN contradicts established literature ^21^ (Fig. 2a). To make comparisons with the TREKING network, which currently includes only kinase-to-kinase connections, we extracted kinase-kinase interactions from the TPS network, yielding a subnetwork of 153 edges (Fig. 2b (bottom left), Supplementary Data S4). Among the 126 kinases predicted by TREKING, 56 (44.4%) kinases were also inferred by the TPS model, demonstrating the ability of these two distinct methodologies to infer the set of kinases activated and play a functional role in barrier regulation. Multiple members of the canonical MAPK cascades were predicted by both TPS and TREKING, including ERKs (MAPK1/ERK2 and MAPK3/ERK1), Jun N-terminal kinases (JNKs; MAPK8/JNK1 and MAPK9/JNK2) and MAPK14/p38α, whose roles in temporal barrier function have been characterized ^13^.

**Fig. 2.**
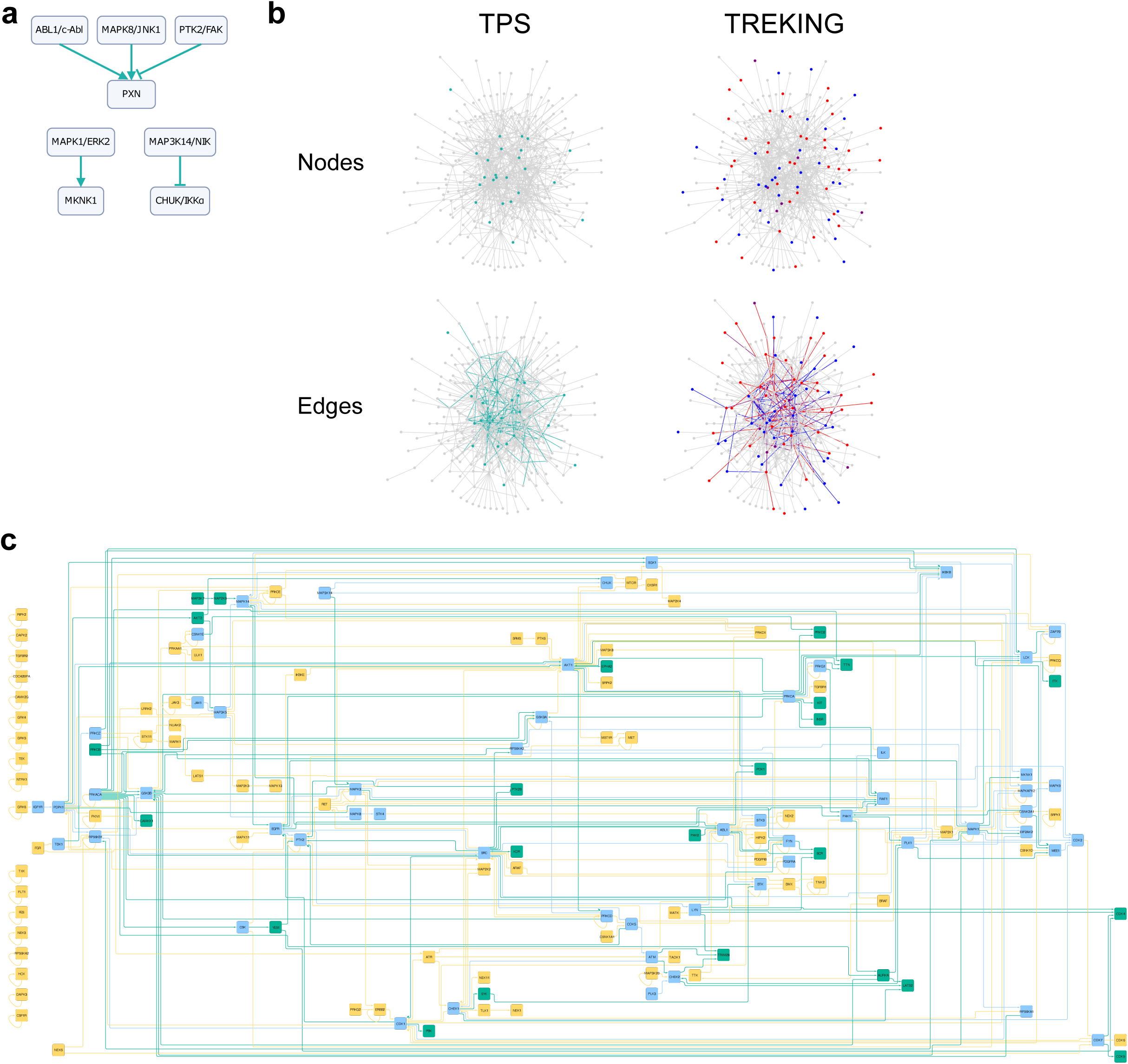
TPS infers common and unique kinases and signaling cascades compared to TREKING models. **a** Signed edges in the summary network generated by TPS. **b** Comparison of kinase phosphosignaling networks built by TPS and TREKING. The background network in gray includes all kinase-kinase phosphorylation interactions from the kinase-substrate phosphorylation database PhosphoSitePlus. Turquoise nodes are the kinases with time series phosphorylation data used to inform the TPS network; turquoise edges are the kinase-kinase phosphorylation interactions inferred by TPS algorithm. Blue and red nodes are the kinases predicted by KiR to have only barrier-weakening and barrier-strengthening functionality, respectively; purple nodes are the switch kinases having both barrier-weakening and barrier-strengthening functionalities but at different time frames. Blue and red edges are the interactions predicted by TREKING that are associated with only barrier-weakening and barrier-strengthening activity, respectively; purple edges are the interactions associated with both barrier-weakening and barrier-strengthening activities. **c** The combined kinase signaling network inferred by TPS and TREKING methodologies. Kinases and kinase-kinase interactions inferred by TPS or TREKING only are in turquoise or moccasin, respectively; kinases and kinase-kinase interactions inferred by both TPS and TREKING are in sky blue.

Additionally, one of the switch kinases, MAPK activated protein kinase 2 (MAPKAPK2/MK2), identified by TREKING as having both barrier-disruptive and barrier-restorative roles, was also predicted by TPS to mediate barrier properties, though TPS does not distinguish between barrier-disruptive or barrier-protective functions.

Among the 153 directed kinase-to-kinase connections in the TPS network, 50 (32.7%) are also present in the TREKING network (Fig. 2b, bottom). Both TPS and TREKING predicted regulation of MAPKAPK2/MK2 by MAPK1/ERK2 and MAPK14/p38α in thrombin-treated endothelial cells (Fig. 2c). TREKING further predicted that the MAPK1/ERK2-MAPKAPK2/MK2 interaction is crucial for barrier disruption, while MAPK14/p38α-MAPKAPK2/MK2 interaction is important for late barrier recovery. Unlike TREKING, TPS does not include phenotypic data in its network reconstruction. Instead, TPS inferred 103 kinase-to-kinase connections not captured by TREKING, including several signaling hubs such as outgoing signals from protein kinase cAMP-activated catalytic subunit alpha (PRKACA), SRC and protein kinase C alpha (PRKCA), and incoming signals to glycogen synthase kinase 3 beta (GSK3B) (Fig. 2c).

TPS’s ability to infer interactions between kinase regulators and their non-kinase substrates makes it a powerful tool for linking upstream kinase signaling to cellular phenotypes, as proteins directly associated with phenotypes are often better characterized and usually not kinases. Despite relying on a sparse input dataset, TPS predicted that cadherin 5 (CDH5), an endothelial adherens junction protein, is regulated by SRC. It also predicted that heat shock protein family B member 1 (HSPB1/HSP27), which directly regulates actin organization in endothelial cells ^22^, is regulated by MAPK kinases MAPK14/p38α and MAPKAPK2/MK2, PRKACA, and protein kinase CGMP-dependent 1 (PRKG1) (Supplementary Data S3); these predictions are supported by experimental validations ^22–24^. Additionally, TPS predicted that PXN, a cytoskeletal protein involved in actin-membrane attachment at focal adhesions, is regulated by Src family kinases FYN and SRC, p21 (RAC1) activated kinase 1 (PAK1), and several MAPK kinases, including MAPK1/ERK2, MAPK3/ERK1, MAPK8/JNK1 and MAPK14/p38α (Supplementary Data S3). These regulatory interactions on PXN and their roles in barrier regulation are consistent with previous studies ^25–27^.

Along with providing a summary network, the TPS pipeline also provides activity windows for each node. However, due to the sparse sampling of the input time series data, TPS could not consistently identify precise activity states (activation, inhibition, or inactivity) for proteins at specific time points, except for those with time course data. This underscores a limitation of TPS in reliably inferring global signaling cascades when working with small input datasets.

We systematically evaluated the accuracy of the predicted barrier-regulatory networks in response to thrombin by comparing the model’s predictions with independent experimental data not used to build the networks (Fig. 3a). For the TPS network, we calculated the cumulative inhibition of each kinase inhibitor on the network (see Methods) as a surrogate for overall perturbation of the barrier-regulatory kinase activity, using the kinase-compound biochemical data ^28^. A cutoff for cumulative inhibition − 25% of the kinases in the TPS network are inhibited by 50% − was set, above which the kinase inhibitor is considered to substantially perturb the network and therefore predicted to alter the barrier permeability. Among the 29 kinase inhibitors in the phenotypic screen ^29^, the model predicts 20 kinase inhibitors to alter the barrier permeability, 18 (90%) of which agree with the measurements (Fig. 3b). We also generated random networks by randomizing both input networks and protein phosphorylation data and ran them through the TPS algorithm (see Methods). The TPS network outperforms random networks in predicting kinase-targeting compounds that modulate barrier permeability (Fig. 3b).

**Fig. 3.**
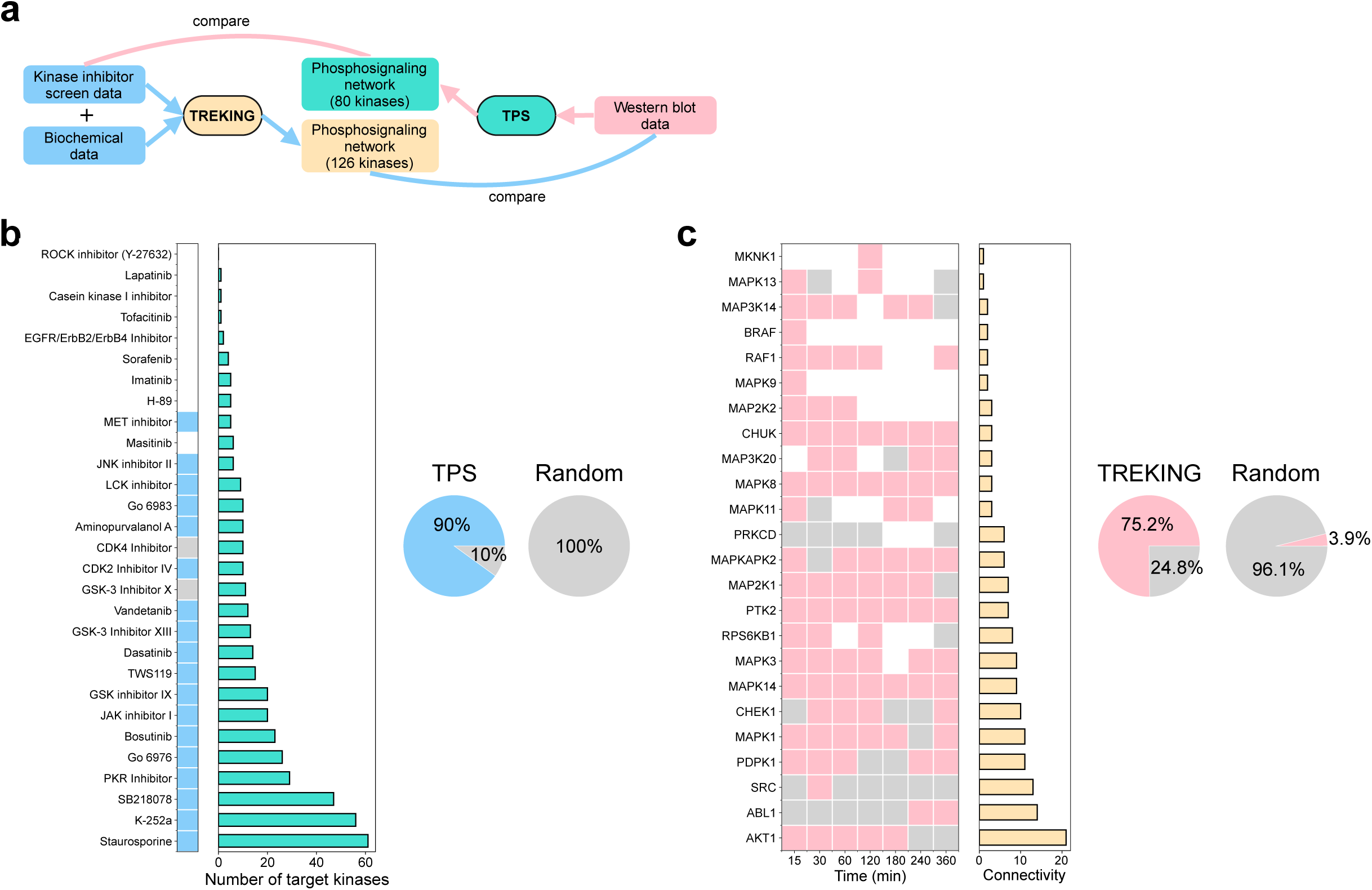
Evaluate the accuracy of model predictions. **a** Schematic showing the types of data used for reconstructing and validating phosphosignaling networks built by TPS and TREKING approaches. **b** Validation on the TPS network using kinase-compound biochemical data ^28^ and kinase inhibitor screen data ^29^. Left: light sky blue indicates that TPS correctly predicted the impact of kinase inhibitors on endothelial barrier permeability, gray indicates incorrect predictions, and white indicates that TPS did not predict substantial inhibition of the network by the kinase inhibitors and no comparison was made with the phenotypic data. Middle: number of target kinases (residual activity < 50%) in the TPS network. Right: overall prediction accuracy of TPS compared to random models on predicting impact of kinase inhibitors on barrier permeability (light sky blue: correct predictions; gray: incorrect predictions). If the model predicts a kinase inhibitor to alter barrier permeability and the barrier permeability in the presence of the kinase inhibitor is different from DMSO-treated control, the model makes correct prediction on the kinase inhibitor; otherwise, the model makes incorrect prediction on the kinase inhibitor. **c** Validation on the TREKING network using western blot data. Left: pink indicates that TREKING correctly predicted the kinase functionality in barrier regulation, gray indicates incorrect predictions, and white indicates that TREKING did not predict the barrier function, and no comparison was made with the western blot data. Middle: kinase connectivity (in/out-degrees of freedom) in the TREKING network. Right: overall prediction accuracy of TREKING compared to random models on predicting the barrier activity of kinases (pink: correct predictions; gray: incorrect predictions). If the model predicts a kinase to regulate barrier permeability and its activity is different from basal level at the corresponding time point, the model makes correct prediction on the kinase barrier activity; otherwise, the model makes incorrect prediction on the kinase barrier activity.

To evaluate the accuracy of the TREKING model for predicting kinase activity, we compared the model predictions of barrier-regulatory kinases with the western blot data used for building the TPS network (Fig. 3a). We first used the TREKING model to generate time-resolved predictions of the activation states of kinases in response to thrombin. To assess the barrier function of a kinase at a particular time point, a 10-minute time window centered at that time point was applied. If any of the signaling pathways with which the kinase is associated are active within the time window, the model predicts the kinase to be functionally active within that time window; otherwise, the model predicts the kinase to be functionally inactive within the time window.

TREKING predicts 129 instances of active barrier function across all time points; 97 (75.2%) differential phosphorylation events were observed by western blot (Fig. 3c). We also generated random networks by randomly sampling temporal profiles for kinases, assigning kinases to random groups, and building phosphosignaling networks based on randomly generated kinase-substrate phosphorylation interactions (see Methods).

Overall, the TREKING network outperforms random models in predicting barrier-regulatory kinases with temporal resolution (Fig. 3c).

In this study, we compared two computational approaches, TPS and TREKING, in reconstructing kinase signaling networks describing phosphosignal propagation within brain endothelial cells after thrombin stimulation. Overall, TPS and TREKING provide different insights into kinase regulatory networks associated with cellular phenotypes.

TPS focuses on the activity of functional proteins, while TREKING focuses on the functionality of kinases altering phenotypes. Since the phosphorylation state of a kinase does not directly imply its function, which is also impacted by its subcellular localization and presence of substrates, discrepancies between the networks reconstructed by these two approaches are expected. Both methodologies are easily generalizable to various biological systems and, in the context of infectious diseases, can deconvolve kinase-mediated signaling mechanisms in host cells or parasites. In the future, integration of TPS and TREKING will facilitate developing more informative network models.

Integrating TPS and TREKING methodologies would allow 1) revealing the functionality of kinases and their inter-connections that alter cellular phenotypes along with their temporal enzymatic activity during changes of cellular phenotypes, and 2) linking functional non-kinase proteins that directly modulate cellular phenotypes to their upstream, functional kinase regulatory pathways that control the activity and function of the proteins.

## Methods

### Cell lines

Primary human brain microvascular endothelial cells (HBMECs; Cell Systems Cat# ACBRI 376) were cultured on rat tail collagen type I (5 µg/cm^2^; Corning Cat# 354236) in HBMEC culture media (Lonza Cat# CC-3202) at 37°C and 5% CO_2_. HBMECs were obtained at passage 3 and used until passage 9.

### Lysate preparation

HBMECs were seeded in 6-well plates (Corning Cat# 353046) at 55,000 cells/well and grown for 4 days with media change every other day. On the day of lysate collection, cells were equilibrated in serum-free culture media for 1 hour and then treated with thrombin at a final concentration of 5 nM for 5, 15, 30, 60, 120, 180, 240, 360 minutes. After the indicated incubation periods, cells were washed twice with ice-cold phosphate buffered saline (PBS) and lysed in sodium dodecyl sulfate (SDS) lysis buffer (50 mM Tris-HCl, 2% SDS, 5% glycerol, 5 mM ethylenediaminetetraacetic acid, 1 mM sodium fluoride, 10 mM β-glycerophosphate, 1 mM phenylmethylsulfonyl fluoride, 1 mM sodium orthovanadate, 1 mM dithiothreitol, supplemented with a cocktail of protease inhibitors (Roche Cat# 4693159001) and phosphatase inhibitors (Sigma-Aldrich Cat# P5726)).

Cell lysates were clarified in filter plates (Pall Cat# 8075) at 2,671×*g* for 30 minutes, after which they were stored at -80°C until use. Cell lysates from three biological replicates were collected.

### Western blot

All the gel electrophoresis was performed using Bolt™ 4-12% Bis-Tris mini protein gels. Proteins were transferred to PVDF membranes using iBlot 2 (Thermo Fisher Scientific Cat# IB21001) or iBlot 3 (Thermo Fisher Scientific Cat# A56727) dry blotting system. Primary antibodies were used at concentrations recommended by the manufacturer (see Supplementary Data S1 for antibody information). Antibody to GAPDH (Cell Signaling Technology Cat# 97166) was used as loading control at 1:2000 dilution. Blots were imaged using Bio-Rad ChemiDoc imaging system and signals were quantified using ImageJ2 (https://imagej.nih.gov/ij/, version 2.3.0). Background correction was done for each band by subtracting background signals nearby the band. The signals from proteins of interest were first normalized to the signals from GAPDH, and then the signals at each time point were normalized to the signals at time zero for the fold change of phosphorylation from basal level. Phosphorylation on activation sites of 25 kinases and 3 non-kinase proteins were evaluated at nine time points (0, 5, 15, 30, 60, 120, 180, 240 and 360 minutes after thrombin treatment). Western blot was performed on cell lysates collected from three biological replicates.

### Reconstruction of TPS network

The input time series data for TPS were collected from western blot experiments. A portion of the data were previously published ^13^. The input network, an undirected subnetwork of the PPI network that connects the phosphorylated proteins to the source node(s), was generated using the Omics Integrator implementation of the Prize-Collecting Steiner Forest algorithm ^17,18^. Time series phosphorylation data, protein prizes and interactome network were required for generating the input network. Protein prizes were obtained using significance scores for each time point relative to the first time point and to the previous time point for each profile. The interactome network was compiled by combining human PPIs from iRefIndex ^30^ (https://irefindex.vib.be//) and kinase-substrate phosphorylation interactions from PhosphoSitePlus ^31^ (https://www.phosphosite.org).

Omics Integrator was run with dummy edge weight of 10, edge reliability of 10, degree penalty of 0, and 100 randomize prize runs. Significance scores for TPS run were computed with paired Student’s t-tests comparing the phosphorylation intensity at each time point with the first time point (“firstscores”) or comparing the phosphorylation intensity at the current time point with the preceding time point (“prevscores”). The threshold for significance scores was set to 0.05, above which measurements are considered non-significant. Proteinase-activated receptor 1 (F2R/PAR1) was designated as the sole source node for TPS run, as it is the predominant thrombin receptor and the only member of PARs mediating thrombin-induced phosphoregulation in endothelial cells ^32,33^. An optional partial model was provided for TPS run, which is a directed kinase-substrate phosphorylation interaction network from PhosphoSitePlus ^31^

(https://www.phosphosite.org).

### Generation of random models

The random TPS summary networks were generated following the TPS pipeline. During each iteration, a random subnetwork and a random partial model were generated by randomly shuffling the nodes in the original networks. Random time series phosphorylation data, random significance scores with respect to the first time point (“firstscores”) or the previous time point (“prevscores”) of the profile were sampled by randomly shuffling the values in the original datasets. A threshold for significance scores was set to 0.05. F2R/PAR1 was designated as the sole source node for generating random summary networks. In total, 100 random summary networks were generated and used to compare with the TPS network.

The random TREKING models were generated following the TREKING pipeline. During each iteration, a random background phosphosignaling network with the same network topology as the background phosphosignaling network used in TREKING was generated using the configuration model function in NetworkX package. All the kinases were shuffled and randomly assigned to each of the nodes in the random network. At each time point, kinases of barrier-weakening or barrier-strengthening functionality were randomly selected from the 300 kinases used for KiR, with the same probabilities as in the TREKING model at the corresponding time point. The random sampling of kinases was performed at all time points corresponding to the TREKING model. Each of the kinases selected at any time point during the time course was randomly assigned to one of the 36 clusters, corresponding to the total number of neurons generated by SOMs.

For each cluster, the phosphosignaling network was reconstructed by searching for the shortest paths between any pair of kinases within the cluster, using the random phosphosignaling network generated for this iteration as the background network. In total, 100 random models were generated and used to compare with the TREKING predictions.

### Visualization

Figs. 1-3 were made in Inkscape (https://inkscape.org, version 1.2.1), a free and open-source vector graphics editor licensed under the GPL. In Fig. 1, the hierarchically-clustered heatmap was generated using seaborn (https://seaborn.pydata.org, version 0.12.0), a Python data visualization library based on Matplotlib; protein-protein interaction network and phosphosignaling networks were visualized using Cytoscape (https://cytoscape.org, version 3.9.1), with layouts generated from yFiles layout algorithms (https://www.yworks.com/products/yfiles-layout-algorithms-for-cytoscape, version 1.1.4); the logo for Omics Integrator was obtained from the website of its original developers (https://fraenkel-nsf.csbi.mit.edu/omicsintegrator/). In Fig. 2, panel (a) was created in BioRender (https://app.biorender.com); phosphosignaling networks in panel (b) were visualized using NetworkX (https://github.com/networkx/networkx, version 2.8.6), a Python package for analyzing complex network structures; phosphosignaling network in panel (c) was visualized using Cytoscape (https://cytoscape.org, version 3.9.1), with its layout generated from yFiles hierarchic layout algorithm (https://www.yworks.com/products/yfiles-layout-algorithms-for-cytoscape, version 1.1.4). In Fig. 3, panel (a) was created in BioRender (https://app.biorender.com); panels (b)-(c) were generated using Matplotlib (https://matplotlib.org, version 3.5.3) in Python.

### Quantification and statistical analysis

The kinase-compound biochemical data ^28^ were used to calculate the cumulative inhibition of kinase inhibitors on the TPS network. For each kinase inhibitor in the screen, its cumulative inhibition was calculated by summing the percentage of inhibition (100 – residual activity) across all the kinases in TPS network (if they are also in the biochemical data).

Student’s t-test was used to evaluate the difference of barrier permeability between kinase inhibitor- and DMSO-treated conditions. The permeability data are from Dankwa *et al.* ^29^ For each kinase inhibitor treatment, both p-value and percent change of normalized area under the curve (AUC) across a 6-hour time window with respect to DMSO-treated condition were calculated to determine if the two sets of data differ from each other. A p-value below 0.05 or all biological replicates reporting a percent change of normalized AUC greater than 10% above/below the DMSO control indicates that the barrier permeability is different between kinase inhibitor- and DMSO-treated conditions. Student’s t-test was also used to evaluate the difference of kinase activity between non-treated and thrombin-treated conditions. At each time point, both p-value and fold change of phosphorylation with respect to non-treated cells were reported to determine if the two sets of data differ. A p-value below 0.05 or all biological replicates reporting a fold change increasing/decreasing by at least 20% compared to non-treated cells indicates that the kinase activity is different between non-treated and thrombin-treated conditions. SciPy statistical package (version 1.9.1) was used to perform the t-tests.

## Data availability

A portion of the data used in this study have been published previously. The remaining data generated in this study is available in the supplementary information.

## Acknowledgments

The authors would like to thank Joseph D. Smith and Selasi Dankwa for helpful discussions. This work was funded by National Institutes of Health (NIH) grants R01 AI148802, R61 HL154250 and R33 HL154250 (to A.K.), and R01 DK041737 and R01 GM109824 (to J.D.A.). F.D.M. is the recipient of a Career Development Award from Seattle Children’s Research Institute. The funders played no role in study design, data collection, analysis and interpretation of data, or the writing of this manuscript.

## Author contributions

**Ling Wei:** Conceptualization, Investigation, Writing – Original Draft, Writing – Review & Editing. **John D. Aitchison:** Funding acquisition, Supervision, Writing – Review & Editing. **Alexis Kaushansky:** Conceptualization, Funding acquisition, Supervision, Writing – Review & Editing. **Fred D. Mast:** Conceptualization, Investigation, Supervision, Writing – Review & Editing.

