## Supplementary Information for "Systems-level reconstruction of kinase phosphosignaling networks regulating endothelial barrier integrity using temporal data"

**This PDF file includes:**

Legends for Supplementary Data S1 to S5

**Other supplementary materials for this manuscript include the following:**

Supplementary Data S1 to S5

**Supplementary Data S1. Antibody information and western blot results on a subset of proteins used to inform TPS network in this study.** This is a Microsoft Excel workbook containing 13 spreadsheets with antibody information and densitometry results for the 11 protein antibody targets (8 kinases, 3 non-kinases) evaluated in this study.

**Supplementary Data S2. The undirected subnetwork generated by the Omics Integrator implementation of the Prize-Collecting Steiner Forest algorithm.** This is a Microsoft Excel workbook containing one spreadsheet.

**Supplementary Data S3. The summary network generated by the TPS algorithm.** The edge types are: A – ProteinA activates ProteinB; I – ProteinA inhibits ProteinB; N – ProteinA regulates ProteinB but the edge sign is unknown. Undirected edges are removed from the summary network. This is a Microsoft Excel workbook containing one spreadsheet.

**Supplementary Data S4. The kinase-kinase edges in the summary network generated by the TPS algorithm.** The edge types are: A – ProteinA activates ProteinB; I – ProteinA inhibits ProteinB; N – ProteinA regulates ProteinB but the edge sign is unknown. Undirected edges are removed from the summary network. This is a Microsoft Excel workbook containing one spreadsheet.

**Supplementary Data S5. Proteins and interactions involved in thrombin signaling through PARs.** Information is from the Reactome pathway – thrombin signaling through proteinase activated receptors (PARs) (stable identifier: R-HSA-456926). This is a Microsoft Excel workbook containing two spreadsheets.
